# JAFFA: High sensitivity transcriptome-focused fusion gene detection

**DOI:** 10.1101/013698

**Authors:** Nadia M. Davidson, Ian J. Majewski, Alicia Oshlack

## Abstract

Genomic instability is a hallmark of cancer and, as such, structural alterations and fusion genes are common events in the cancer landscape. RNA sequencing (RNA-Seq) is a powerful method for profiling cancers, but current methods for identifying fusion genes are optimized for short reads. JAFFA (https://code.google.com/p/jaffa-project/) is a sensitive fusion detection method that clearly out-performs other methods with reads of 100bp or greater. JAFFA compares a cancer transcriptome to the reference transcriptome, rather than the genome, where the cancer transcriptome is inferred using long reads directly or by *de novo* assembling short reads.

## Background

Chromosomal rearrangements have the potential to alter gene function in many different ways; for example, they may produce chimeric fusion proteins that gain new functionality, or place a gene under the control of alternative regulatory elements [1, 2]. Fusion genes including BCR-ABL, PML-RAR and EML4-ALK have become targets for therapy in cancer, and as a result there is great interest in defining the full complement of oncogenic fusion genes.

Next generation sequencing of RNA (RNA-Seq) has greatly accelerated the discovery of novel fusion genes in cancer [3–5]. However, while a large number of tools have been presented to identify fusion event using RNA-Seq [5–9], practical use of fusion finding tools is often hampered by either a high false detection rate or low sensitivity [10, 11]. Many fusion detection methods identify transcriptional breakpoints by splitting short reads into even shorter segments and then aligning these segments to the genome [5, 12]. Short read sequences have lower alignment specificity particularly in the presence of SNPs, sequencing errors and repeat regions. Incorrect mapping of these short read fragments has the potential to lead to false predictions. To overcome this, algorithms look for supporting information, such as neighbouring reads, or read pairs, that cover the same breakpoint. This strategy can be effective at controlling the false discovery rate, but often requires restrictive filtering that may limit sensitivity.

Another limitation of many fusion finding algorithms is that they have been built and tested using reads shorter than 100bp. Sequencing reads are becoming longer, with 100bp paired-end reads now standard for many applications, and read lengths promise to grow in the coming years. The Mi-Seq and PacBio platforms already produce reads of several hundred and several thousand bases respectively. It is not clear how current fusion finding algorithms will perform on long read data. For example, many will not work on long single-end data, because they require paired-end reads.

In this study we outline a new method for detecting fusion genes that can be applied to any read length, single or paired-end. A critical and unique feature of our method is that rather than comparing a tumour transcriptome to the reference genome we compare it to the reference transcriptome. There are several advantages in alignment to the transcriptome rather than genome; the complexity of splice site alignment, which can be error prone [13, 14], is avoided as the transcriptome only includes exonic sequence; identifying fusion transcripts from those alignments is simplified because we do not need to check if the break can be explained by splicing; and finally, the reference transcriptome consists of less sequence than the reference genome, allowing for slower, but more accurate alignment algorithms to be used, such as BLAT [15]. Critically, BLAT works well over a range of reads lengths, whereas mapping algorithms used by other fusion finders are optimized for short reads. For example, bowtie [16], the recommended aligner for TopHat-Fusion [6], will not map reads longer than 1024 bases.

Our new method, called JAFFA, is designed for detecting fusion in RNA-seq data with contemporary read lengths. Fusions may be identified using reads from 100bp up to full-length transcripts. Reads shorted than 100bp can be analysed effectively by assembling them *de novo* into contigs of 100bp or longer – a step which is performed by JAFFA. Hence, JAFFA is a complete pipeline; it uses *de novo* assembly or raw reads directly to align to a reference transcriptome and outputs candidate fusions along with associated information such as the position of the break in the genome, a prediction of reading frame, read support metrics and whether the fusion is present in the Mitelman database [17]. JAFFA also reports the sequence of the fusion read or assembled contig. JAFFA is built using the Bpipe platform [18] and takes advantages of features such as modularity of the pipeline stages, running numerous samples in parallel, and integration with computing clusters. JAFFA is therefore a highly effective tool for large RNA-Seq studies involving multiple datasets and samples. The idea behind JAFFA has already been used to successfully identify fusions in lung cancer [19].

We validated JAFFA on a range of data with different read-lengths, including 50bp, 75bp, 100bp paired-end reads and ultra-long PacBio reads [20, 21]. We used RNA-Seq from breast cancer cell lines [22], glioma tumours [23], as well as simulation and found JAFFA has a low false discovery rate without compromising on sensitivity. JAFFA may be run in three defined modes: assembling short reads (shorter than 60bp), using long reads directly (100bp or greater), or a hybrid approach that both assembles and processes unmapped reads (between 60bp and 100bp). We performed a detailed comparison to established methods and found that JAFFA consistently gave the best performance on contemporary data with reads longer than 50bp. On 100bp datasets, JAFFA’s computational requirements were comparable to those of other fusion finding tools.

## Results and discussion

### The JAFFA method

JAFFA is a multi-step pipeline that takes raw RNA-Seq reads and outputs a set of candidate fusion genes along with their cDNA breakpoint sequences. JAFFA runs in three modes: (i) “Assembly” mode assembles short reads into transcripts prior to fusion detection (ii) “Direct” mode uses RNA-Seq reads directly, rather than assembled contigs, by first selecting reads that do not map to known transcripts, or (iii) “Hybrid” mode both assembles transcripts and supplements the list of assembled contigs with reads that do not map to either the reference transcriptome or the assembly. The appropriate mode to use depends on the read length (Additional File 1: Supplementary Figure 1). By default, JAFFA requires 30 bases of flanking sequence either side of the breakpoint. For reads shorter than 60bp, the flanking sequence would be too short to accurately and efficiently align using BLAT, so the Assembly mode must be used. For reads 60-99bp long, Hybrid mode is used, while for reads 100bp and over there is no advantage in performing a *de novo* assembly so the Direct mode is used. When *de novo* assembly is performed, Oases [24] is used. We found Oases gave superior sensitivity compared with other assemblers (Additional File 1: Supplementary Material 1, Additional File 2). *De novo* assembly is well known to producing a high fraction of false chimeras [25, 26] and we found an effective method to control for these by checking the amount of sequence shared by fusion partner genes at the breakpoint (Additional File 1: Supplementary Material 1, Additional File 1: Supplementary figure 2).

JAFFA is based on the idea of comparing a sequenced transcriptome against a reference transcriptome. As a default, JAFFA uses transcripts from GENCODE [27] as a reference. For all JAFFA modes, reads aligning to intronic or intergenic regions are first removed to improve computational performance (step 1 in Figure 1, see Materials and methods). Sequences are then converted into a common form – tumour sequences – consisting of either assembled contigs or the reads themselves. These sequences are processed by a core set of fusion-finding steps (steps 2-6 in Figure 1). First, sequences are aligned to a reference transcriptome and those that align to multiple genes are selected. Second, read support is determined. Third, putative candidates are aligned to the genome to check the genomic position of breakpoints. Finally, JAFFA calculates characteristics of each fusion and uses this to prioritize candidates for validation. Each of these pipeline steps is described in detail in Materials and Methods.

**Figure 1.**
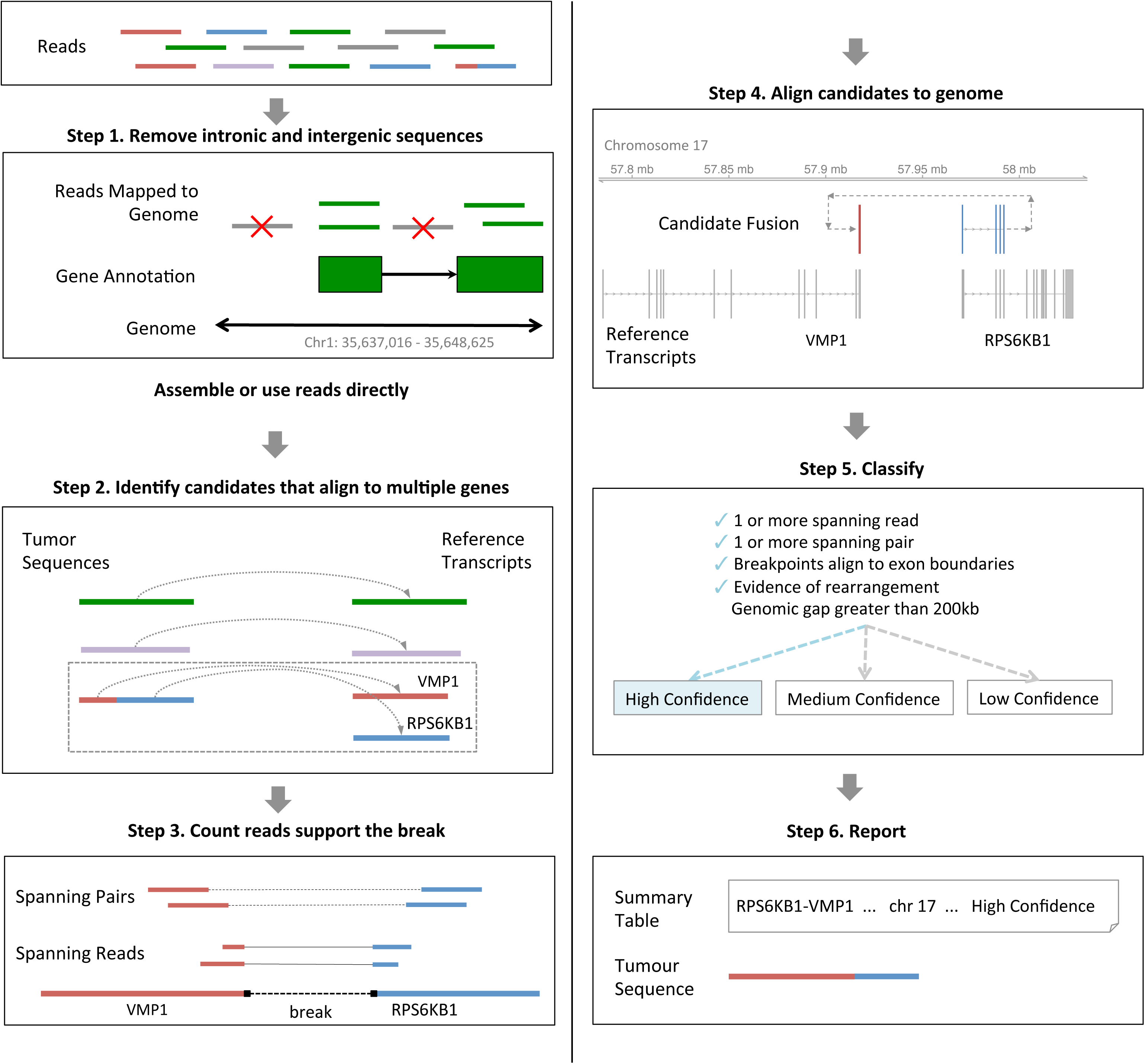
The JAFFA Assembly pipeline. An example of the JAFFA pipeline is demonstrated in detail using the RPS6KB1-VMP1 fusion from the MCF-7 breast cancer cell line dataset (Step 1.) RNA-Seq reads are first filtered to remove intronic and intergenic reads. 50bp reads would then be assembled into contigs using Oases. For longer reads, this step is not necessary. (Step 2.) The resulting tumour sequences are then aligned to the reference transcriptome and those that align to multiple genes are selected. These contigs make up a set of initial candidate fusions. Next, (Step 3.) the pipeline counts the number of reads and read pairs that span the breakpoint. (Step 4.) Candidates are then aligned to the human genome. Genomic coordinates of the breakpoints are determined and (Step 5.) further selection and candidate classification is carried out using quantities such as genomic gap size, supporting reads and alignment of breakpoints to exon-exon boundaries. (Step 6.) A final list of candidates is reported along with their sequence.

Most fusion genes originate from a genomic rearrangement with breakpoints in intronic DNA. We found empirically that transcriptional breakpoints aligning to exon-exon boundaries were more indicative of a true fusion than the number of reads supporting the breakpoint, and have incorporated this into our ranking system. Genes with breakpoints aligning to exon-exon boundaries are classified as either “High Confidence” or “Medium Confidence”. These two categories are distinguished by either the presence (“High Confidence”) or absence (“Medium Confidence”) of both spanning reads and spanning pairs. Spanning reads have the fusion breakpoint sequenced within the read. Spanning pairs lie on opposite sides of the breakpoint (Step 3 in Figure 1). For single-end data, only “Medium Confidence” is reported because spanning pairs are not calculated. Unlike other fusion finding algorithms, such as deFuse and TopHat-Fusion, which apply a threshold on the number of supporting reads to ensure the false discovery rate is controlled, JAFFA can detect fusions with a single read, without compromising the false discovery rate. Fusions with spanning pairs, but without transcriptional breakpoints aligning to exon boundaries are classified as “Low Confidence”. For “LowConfidence” fusions we require two spanning reads so that chimeric artifacts produced during library preparation are removed. Fusions without spanning pairs or breakpoints aligning to exon boundaries are discarded. Finally, JAFFA flags a fourth class of candidates “Potential Regular Transcript”, which appear to be novel transcripts between adjacent genes [28]. We identify these by a genomic gap between the breakpoints of less than 200kb and no evidence for genomic rearrangement. Because these candidates are likely to be caused by read-through transcription [29], they are excluded from the default reporting of our software. For candidates within a class, we rank by the sum of spanning reads and spanning pairs. When read support is equal, we rank on the genomic gap size, with smaller gaps ranked higher. We did this because we found empirically that true positives were often intrachromosomal and localized (Additional file 1: Supplementary figure 3).

Because JAFFA is a pipeline rather than a standalone software tool, many of its stages rely on external software. The choice of these programs, the reference annotation and genome can be easily customized. In JAFFA, bash and R scripts are used to steer each step, and the pipeline is implemented using the Bpipe platform [18]. Bpipe handles parallelization, restarting from midway through the pipeline and error reporting, and is convenient for analyses involving a large number of samples. In Material and Methods, we describe each stage of JAFFA version 1.04 in more detail along with the software choices used during validation. JAFFA is open source and available for download from https://code.google.com/p/jaffa-project/

### Datasets and competing tools used to assess JAFFA

JAFFA’s sensitivity and false discovery rate were evaluated on several datasets. Firstly, we used simulated data provided by FusionMap [8] to assess JAFFA’s power. The *FusionMap dataset* consisted of 57 thousand 75bp pair-end RNA-Seq reads. 50 fusion events were simulated, with a range of coverage levels. However, background reads from non-fusion genes were absent. Therefore we simulated a second dataset to validate JAFFA’s false discovery rate by generating 20 million, 100bp paired-end RNA-Seq reads without fusion events – the *BEERS dataset*. The simulation was performed using BEERS [30] with default parameters.

Next, we assessed JAFFA’s performance using RNA-Seq of several breast cancer cell lines, for which numerous fusions have previously been reported and validated. We did this for a range of read lengths: firstly, we ran the Assembly mode on 50bp paired-end reads from Edgren et al. [22]. The *Edgren datasets* contained between 14 and 42 million, 50bp paired-end reads of each of the BT-474, SK-BR-3, KPL-4 and MCF-7 cell lines. Next we used the *ENCODE dataset* containing 40 million 100bp paired-end reads of the MCF-7 cell line to assess JAFFA’s Direct mode [21]. We also assessed the Direct mode on an MCF-7 transcriptional profiling dataset provided by PacBio [20]. The *PacBio dataset* consisted of 44,531 non-redundant consensus sequences. In the BT-474, SK-BR-3, KPL-4 and MCF-7 cell lines, used in the *Edgren dataset*, a total of 99 fusions have previously been validated (Additional file 3) [22, 31–34]. We used these fusions as our set of true positives. It is worth noting that not all previously published fusions are identified in all datasets. This is likely not only because of limitations by fusion detection tools, but also because of differences in sequencing methodology, depth and because of variation in cell line preparations from different laboratories. The concordance between different datasets of the MCF-7 cell line is provided in Additional file 1: Supplementary figure 4.

Finally, we ran JAFFA on 100bp paired-end RNA-Seq from a large glioma study [23]. From the full dataset of 272 samples, we selected a subset of 13 samples to form our *glioma* validation dataset Each of these samples contained 2 or more validated inframe fusions, with 31 true positives in total (Additional File 4).

We compared JAFFA to four of the most widely used fusion detection methods; TopHat-Fusion [6], SOAPfuse [35], DeFuse [7] and FusionCatcher [36]. This choice was based on the results from several studies [6,9,10, 22], along with our own assessment of a broader selection of tools using the Edgren and FusionMap datasets (Summarized in Additional file 1: Supplementary table 1). TopHat-Fusion and DeFuse are older fusion finding programs, but are used broadly. FusionCatcher and SOAPfuse have been released more recently and promise superior performance over existing tools.

### JAFFA shows good sensitivity and a low false discovery rate on simulated data

The performance of JAFFA was first assessed using the 75bp paired-end reads of the FusionMap simulation. JAFFA was run using all three modes: Assembly, Direct, and Hybrid (Table 1). JAFFA’s Assembly mode reported 39 out of 50 true positives (78% sensitivity). For the Direct mode this value was lower, at 34 (68% sensitivity). Finally, the Hybrid approach reported more true positives than any other tool (44 out of 50, 88% sensitivity), indicating that even with reads as short as 75bp, searching for fusions amongst reads in addition to assembly, improves sensitivity. For all JAFFA modes, true positives were reported as either “High Confidence” or “Medium Confidence”. The majority of missed true positives had low read coverage. In contrast to the previous finding of a high false positive rate with the FusionMap dataset (Carrara et al. [10, 11], Additional File 1: Supplementary Table 1A), we found that JAFFA, TopHat-Fusion, FusionCatcher, SOAPfuse and deFuse all had very high specificity, with only SOAPfuse reporting one false positive (Table 1).

**Table 1.** A comparison of fusion detection performance on simulated RNA-Seq. We ran all three modes of JAFFA in addition to SOAPfuse, TopHat-Fusion, deFuse and FusionCatcher on a simulation set of 57 thousand 75bp RNA-Seq read pairs provided with FusionMap. JAFFA had the highest sensitivity when run in Hybrid mode, identifying 44 out of 50 possible fusion events. For all JAFFA modes, no false positives were reported. In parenthesis we show the value at each of JAFFA’s classifications levels: (high / medium / low) confidence.

|  | True Positives | Sensitivity | False Positives |
| --- | --- | --- | --- |
| JAFFA - Hybrid | 44 (32/12/0) | 88% | 0 |
| JAFFA - Assembly | 39 (28/11/0) | 78% | 0 |
| SOAPfuse | 37 | 74% | 1 |
| JAFFA - Direct | 34 (32/2/0) | 68% | 0 |
| deFuse | 34 | 68% | 0 |
| TopHat-Fusion | 27 | 54% | 0 |
| FusionCatcher | Unable to run on a low number of reads |  |  |

Because the FusionMap simulation contained no background reads, we assessed JAFFA’s false positive rate further with a simulation containing no fusions, but with transcriptional run-through events, the BEERS dataset. On this dataset JAFFA reports no false positives with a rank of “High Confidence” or “Medium Confidence” in all modes. However, the Assembly and Hybrid modes did report 23 “Low Confidence” false positives. These false positives were misassembled because of sequence homology along with sequencing errors, SNPs and indels. However, because exon-exon alignment was not preserved, they were ranked as “Low Confidence”. Across all datasets we tested, JAFFA almost always classified true positives as either “High Confidence” or “Medium Confidence”. Therefore, in practice, we advise that “Low Confidence” candidates be rejected, unless there is other independent information to support them. JAFFA’s Direct mode, which is the nominal mode for the BEERS 100bp reads, did not report false positives at any classification level. TopHat-Fusion also did not report any false positives on the BEERS dataset. SOAPfuse reported 111 candidate fusions and FusionCatcher 79, however in both cases, the tools flagged these false positives as transcriptional run-through events. DeFuse reported 215 false positives, of which 156 where classified as run-through transcription.

### JAFFA has excellent performance across a range of read lengths on cancer RNA sequencing

#### Short reads (50bp)

On the Edgren dataset, SOAPfuse reported the highest number of true positives, 41, with other tools reporting between 27 and 35 (Table 2A). Of the 40 validated fusions previously published for the Edgren dataset [22, 34], 37 were rediscovered by at least one of the tools tested. In addition, 8 fusions that had been validated in other datasets [31–33] of the same cell lines were reported by at least one tool. Of the total 48 true positives, JAFFA missed 20, predominantly as a result of failing to be assembled (e.g. Additional File 2).

**Table 2.** A comparison of fusion detection performance on cancer RNA-Seq. (A) The Edgren dataset, consisting of between 7 and 21 million 50bp read pairs of the BT-474, SK-BR-3, KPL-4 and MCF-7 cell lines. Using a list of 99 validated fusions in these cell lines, we compared the predictions of JAFFA to TopHat-Fusion, SOAPfuse, deFuse and FusionCatcher. In total, 48 true positives have been reported for this dataset. Predictions not in the list of validated fusions, but involving one of the partner genes in the list of validated fusions, or fusions that were predicted by three or more tools are designated as probable true positives. (B) We compare JAFFA against alternative tools on the ENCODE dataset which consists of 20 million read pairs of MCF-7. Combing the results of all tools, 30 true positives were observed. JAFFA reports more true positives than the other methods. (C) JAFFA’s high sensitivity is also seen on 100bp paired-end dataset from 13 glioma samples for which 31 true positives are known. The samples range from 15 to 35 million read pairs.

**A) Edgren breast cancer cell line dataset**
|  | True Positives | Probable True Positives | Other Positives |
| --- | --- | --- | --- |
| SOAPfuse | 41 | 1 | 18 |
| TophatFusion | 35 | 8 | 218 |
| deFuse | 29 | 4 | 43 |
| JAFFA - Assembly | 28 (24/3/1) | 1 (0/0/1) | 13 (2/3/8) |
| FusionCatcher | 27 | 1 | 3 |

B) ENCODE breast cancer cell line dataset
|  | True Positives | Probable True Positives | Other Positives |
| --- | --- | --- | --- |
| JAFFA – Direct | 27 (19/8/0) | 9 (3/6/0) | 111 (6/101/4) |
| SOAPfuse | 22 | 2 | 46 |
| deFuse | 16 | 14 | 90 |
| FusionCatcher | 16 | 2 | 14 |
| TopHat-Fusion | 12 | 4 | 28 |

C) Gliomas dataset
|  | True Positives | Probable True Positives | Other Positives |
| --- | --- | --- | --- |
| JAFFA – Direct | 30 (30/0/0) | 96 (47/44/5) | 3888<br>(149/3208/530) |
| deFuse | 29 | 53 | 616 |
| TopHat-Fusion | 29 | 25 | 252 |
| FusionCatcher | 28 | 42 | 146 |
| SOAPfuse | 22 | 41 | 236 |

In addition to the true positives, all tools reported a number of additional candidates. A subset of these are likely to be novel true positives, and we attempted to distinguish these from other reported candidates using either of the following criteria: *i)* candidates reported by three or more tools, after excluding those marked as run-through transcription (Additional File 1: Supplementary figure 5); or *ii)* candidates where one of the partner genes is implicated in a true positive fusion in the same sample. For example, an unconfirmed candidate, SULF2-ZNF217 was identified by JAFFA in the MCF-7 cell lines. Because MCF-7 harbours validated fusions involving SULF2 (SULF2 partnered with ARFGEF2, NCOA3 and PRICKLE2), SULF2-ZNF217 was counted as a probable true positive (Table 2A). These so called “promiscuous fusion gene partners” were also observed to occur within the same sample (the MCF-7 and BT-474 cell lines) by Kangaspeska et al. [34]. Kangaspeska et al. noted that some promiscuous fusion gene partners were amplified and speculate the mechanism for multi-fusion formation may involve breakage-fusions-bridge cycles where the breakage repeatedly occurs within the same gene.

The number of other reported positives that were neither true positives, nor probable true positives varied substantially between each tool, from 3 (FusionCatcher) to 218 (TopHat-Fusion). The absolute number of other reported positives is often not as informative as assessing the ranking of positives, which we did using an ROC style plot (Figure 2A). DeFuse and TopHat-Fusion each provided a probability value to rank candidates on. For other tools, we ranked using the output information that maximized the area under the ROC curve. For both FusionCatcher and SOAPfuse this was the number of spanning reads. Probable true positives were excluded from the plot. SOAPfuse, FusionCatcher and JAFFA ranked most known fusions high, however SOAPfuse achieved far greater sensitivity that all other tools without compromising on false discovery rate.

**Figure 2.**
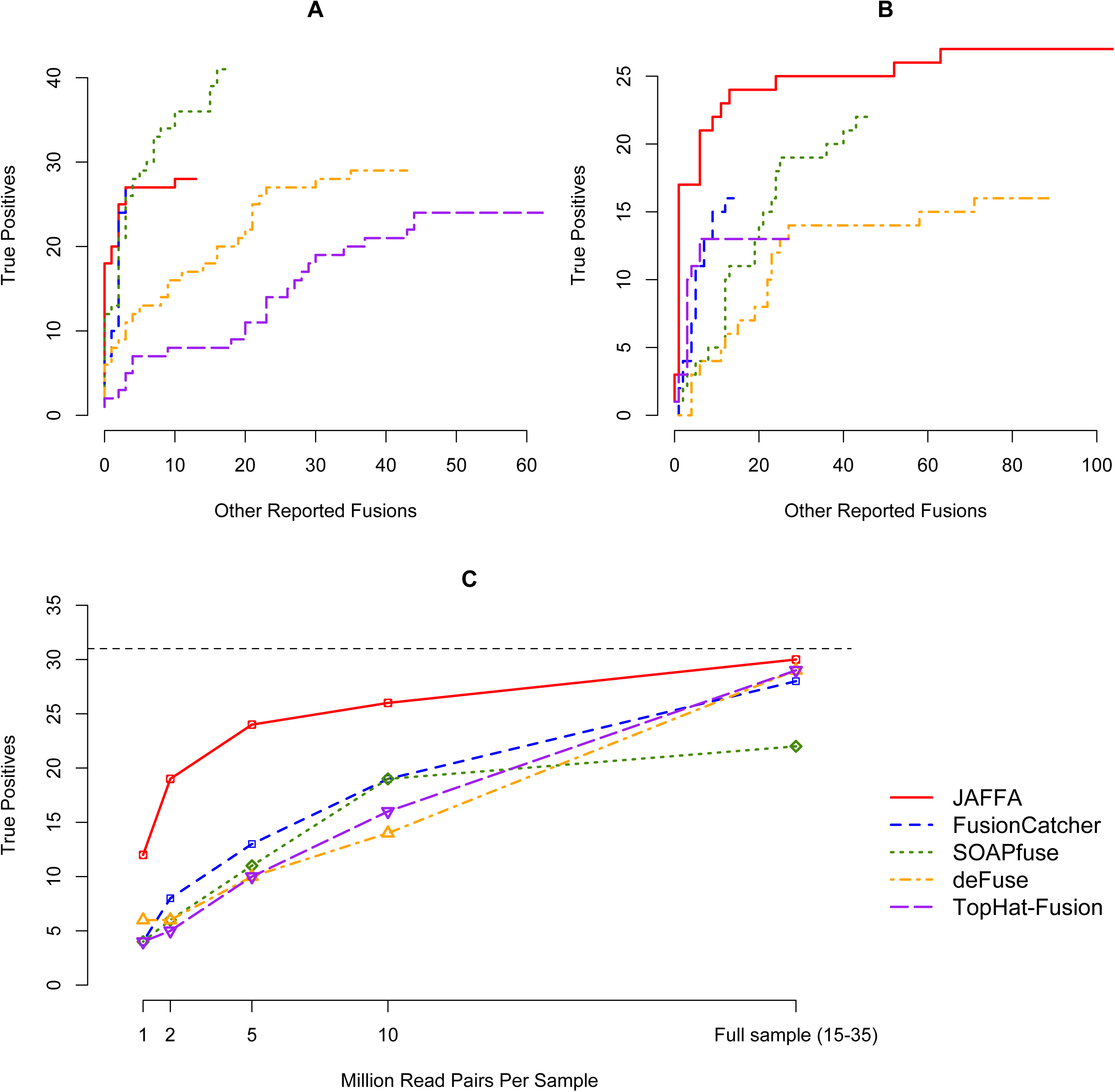
Performance of JAFFA and four other tools on cancer RNA-Seq. (A) An ROC-style curve for the ranking of candidate fusions in the Edgren dataset. The Edgren dataset consists of between 7 and 21 million 50bp read pairs of the BT-474, SK-BR-3, KPL-4 and MCF-7 cell lines. The number of true positives are plotted against the number of other reported positives from a ranked list of fusion candidates. Probable true positives (see text for detail) are removed. Higher curves indicate a better ranking of the true positives. For each fusion detection tool, we ranked the candidates using the tools own scoring system, or if absent, the supporting data that maximized the area under the curve. SOAPfuse ranked true positives higher than other tools, followed by FusionCatcher and JAFFA. (B) On long read data - the ENCODE dataset consisting of 20 million 100bp read pairs of the MCF-7 cell line - JAFFA ranks true positives higher than any other tool. (C) JAFFA’s sensitivity is confirmed on a second long read dataset – 13 glioma samples with read depths varying from 15-35 million 100bp read-pairs. JAFFA identifies 30 of the 31 true positives (total true positives are indicated by the dashed line). Downsampling the data to mimic smaller read depths indicates that JAFFA has similar sensitivity with 2 million read pairs per samples as other tools on 10 million read pairs per sample.

All tools had similar computational performance, with the exception of TopHat-Fusions taking longer to run (27 hours on a single core of a modern computing cluster compared to under 11 hours for all others). Unlike the other tools, JAFFA’s RAM utilisation was not constant, but scaled with the input reads due to the *de novo* assembly (Additional File 1: Supplementary figure 6A and 6B).

#### Long reads (100bp)

JAFFA’s Direct mode, which is suitable for reads of 100bp and longer was assessed on the ENCODE MCF-7 data (Table 2B, Figure 2B). JAFFA reported the highest number of true positives (27) of the fusion detection tools and a large number of probable true positives (9), however JAFFA also reported the highest number of other positives (111). These were largely classified as “Medium Confidence” (92% of candidates) and supported by only a single read (91%). 32% of the other reported positives were intrachromosomal, and 19% had a genomic gap of less than 3Mb. The proportion of local rearrangements were consistent with fusions in the Mitelman database [17] (Additional file 1: Supplementary figure 3). We note that JAFFA’s predictions are inconsistent with the false positives reported for the BEERS simulation. Those false positives were reported only for the JAFFA modes involving assembly, were classified as “Low Confidence” and had a more random genomic distribution (Additional file 1: Supplementary figure 3). Another interesting possibility, is that the unknown positives are trans-splicing events, such as those found in normal tissue [37, 38]. These are often also localized [39, 40]. Despite the larger number of unknown positives, JAFFA out-performed all other tools in its ability to rank true positives before other positives (Figure 2B). Again, probable true positives were excluded from the ROC curve. Finally, we compared JAFFA’s Direct mode against the Hybrid and Assembly modes (Additional File 1: Supplementary Figure 7, Supplementary Table 2), which confirm that there is no advantage in performing an assembly for longer reads (>100bp). On the contrary, assembly requires substantially more computational resources (Additional File 1: Supplementary Figure 6C and 6D).

As a validation of the superior performance of JAFFA with 100bp reads, we assessed a second dataset consisting of 13 glioma samples with 31 validated fusions. JAFFA reported the highest number of true positives (30 out of 31) and the highest number of probable true positives (95) (Table 2C). Many of the probable true positives can be explained as out-of-frame fusions that were not validated by Bao et al., as only inframe fusions were followed-up for validation. TopHat-Fusion and DeFuse reported the equal second highest number of true positives (29), however, we note that the fusions validated by Bao et al. were first identified as the intersection of candidates reported by these two tools. We have attempted to avoid the bias that favours TopHat-Fusion and DeFuse by downsampling the dataset to depths of 1,2,5 and 10 million read pairs per sample. Across the range of read depths, JAFFA had significantly higher sensitivity in all cases (Figure 2C), while consistently ranking those true positives highly (Additional File 1: Supplementary Figure 8). For example, we found that JAFFA analyzing just 2 million read pairs achieved the same sensitivity as all other tools on 10 million read pairs, without compromising the false discovery rate (Additional File 1: Supplementary Figure 9). The sensitivity of JAFFA comes from its ability to reliably call fusions with very low coverage. For example, three of the true positives detected exclusively by JAFFA on the 2 million pair dataset, had just a single read supporting them. This high sensitivity may allow fusions to be identified in samples with low tumour purity or in samples in which a particular fusion is only present in a proportion of tumour cells. The other positives reported by JAFFA, of which there were approximately 300 per sample, displayed similar characteristic to those in the ENCODE dataset, such as a high number of localized rearrangements (Additional File 1: Supplementary Figure 3).

On 100bp reads, all tools were comparable in terms of computational performance (Additional File 1: Supplementary Figure 6E and 6F). On the ENCODE dataset, containing 20 million read-pairs, the fusion finding programs took from 7-20 hours on a single core and 6-13 GB of memory. JAFFA required 16 hours and 8 GB of RAM. On the gliomas dataset, 13 samples ranging from 15-35 million read-pairs were run in parallel. The fusion finding tools required 13-50 hours and 6-13 GB of RAM. JAFFA took 23 hours and 11 GB of RAM. Across the Edgren, ENCODE and gliomas datasets, FusionCatcher was consistently the fastest and SOAPfuse consistently used the least memory.

#### Ultra-long reads and pre-assembled transcriptomes

Read lengths are increasing, and technologies such as Ion Torrent, MiSeq and PacBio can already produce reads from several hundred bases up to several kilobases. JAFFA is intrinsically designed for the analysis of such data, because it is based on the idea of comparing transcriptomes. By contrast, it is unclear how well other short read tools work on these data. For example, SOAPfuse, FusionCatcher and deFuse require paired-end reads. TopHat-Fusion could not be run with its recommended aligner, bowtie, because bowtie only aligns reads 1024bp and shorter, whereas the PacBio dataset has an average sequence length of 1,929 bp. Bowtie2, which aligns longer reads, may also be used with TopHatFusion, but we were not able to run it successful.

To assess the performance of JAFFA on long reads, we ran the Direct mode on the PacBio dataset. Compared to PacBio’s own fusion predictions, released with the data [20] (software unavailable), JAFFA reported a similar number of true or probable true positives (17 compared to 18), but fewer other positives (5 compared to 64). The 5 unknown positives reported by JAFFA, were also predicted by PacBio. One of these was also predicted by JAFFA in the ENCODE dataset. These results indicate that JAFFA has excellent specificity on ultra-long reads, while still achieving sensitivity similar to tools purpose built for such reads.

## Optimal choice of read layout and length

Using the ENCODE dataset, we next addressed the questions of whether paired-end reads perform better than single-end reads, and whether there is any advantage in using 100bp reads over 50bp. This question aims to inform experimental design when the sequencing costs of 100bp, 50bp, single-end and paired-end are similar for a given number of total bases sequenced. The ENCODE dataset has 100bp paired-end reads, and was used to create pseudo single-end reads, by selecting one read from each pair, and pseudo 50bp reads, by trimming off the final 50 bases of each read. JAFFA’s Assembly mode was run on the 50bp reads and the Direct mode was run on the 100bp reads. Each dataset was created with 4 billion sequenced bases – i.e. 20 million 100bp pairs, 40 million 100bp single-end reads, 40 million 50bp pairs and 80 million 50bp single-end reads. Note that the 20 million 100bp pairs were the same dataset used for the 100bp validation presented earlier in this manuscript.

When considering each combination of read layout, length and fusion finding algorithm, we found that JAFFA with 100bp paired-end reads produced the highest number of true positives, with a total of 27 (Figure 3A). However, defuse, SOAPfuse and TopHat-Fusion reported a similar number of true positives on 50bp paired-end reads with 26, 24 and 24 respectively. To determine if these tools were effective at separating the true positives from other preditions, we used an ROC-style curve (Figure 3B). For each tool we show the combination of read length and layout that maximized the ROC performance. For SOAPfuse, deFuse and TopHat-Fusion, this was 50bp paired-end reads and for JAFFA and FusionCatcher, 100bp paired-end reads. JAFFA on 100bp paired-end reads not only reported the highest number of true positives, but provided the best ranking of those true positives (Figure 3B). This trend held across a range of sequencing depths (250 million and 1 billion sequenced bases, Additional file 1: Supplementary figures 10 and 11).

**Figure 3.**
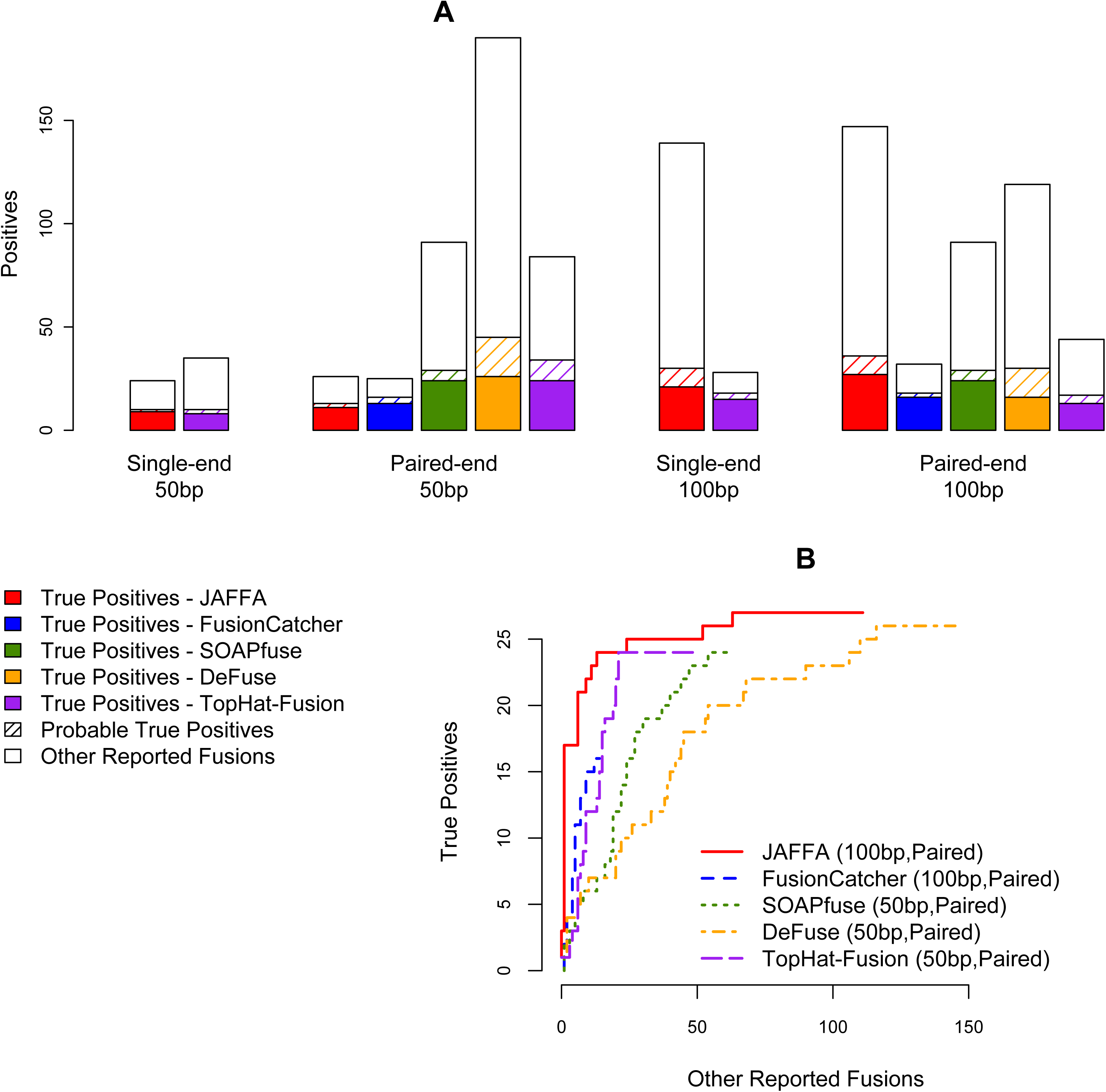
Performance of JAFFA and four other tools for different read lengths and layouts. We compared the performance of JAFFA, FusionCatcher, SOAPfuse, deFuse and TopHat-Fusion on the ENCODE dataset of the MCF-7 cell line, trimmed to emulate four different read configurations: single-end 50bp (80 million reads), paired-end 50bp (40 million read pairs), single-end 100bp (40 million reads) and paired-end 100bp (20 million read pairs). In each case, the total number of bases sequence was 4 billion. Only JAFFA and TopHat-Fusion could process single-end data. (A) Most true positives were reported with JAFFA on 100bp paired-end reads followed by deFuse on 50bp paired-end reads. (B) For each tool we compared the ranking of fusions, by selecting the read length and layout that maximised ROC performance. JAFFA on 100bp reads ranked true positives higher than any other combination.

Taken together with the results from the simulation, Edgren and glioma datasets, we recommend that datasets with 50bp paired-end reads be analysed with SOAPfuse, whereas reads longer than 50bp or single-end reads should be analysed with JAFFA. When considering how to design an experiment to detect fusion genes, it appears that optimal performance is obtained with 100bp paired-end sequencing followed by analysis using JAFFA.

## Conclusions

We have presented JAFFA, a method for the discovery of fusion genes in cancer transcriptomes by comparing them to a reference transcriptome. The cancer transcriptome is either a set of contigs created by *de novo* assembly of short reads or the reads themselves for longer read sequencing. Therefore one major advantage of JAFFA over previous methods is that it detects fusions using RNA-seq reads of any length, with either single or paired-end reads. JAFFA also provides a simple and effective method of ranking fusions based on read support and exon-exon boundary alignment. This approach means that we avoid restrictive filtering that may reduce sensitivity.

A limitation of our approach is that JAFFA is not sensitive to fusion genes incorporating intronic or intragenic sequence, because the reference includes only exonic sequence. Moreover, JAFFA down ranks fusions when the breakpoint occurs within an exon, rather than at the boundary. In this case the fusion is ranked as “Low Confidence”. These two classes of fusions are rare [41, 42] and we argue that on balance, the overall improvement in sensitivity and ranking outweighs the potential for these fusion types to be missed. In addition, because JAFFA reports whether a fusion is found in the Mitelman database, <u>fusions classified as</u> “Low Confidence” <u>that are recurrent in cancer</u> remain identifiable to the user.

The validation of JAFFA on simulation and RNA sequencing of cancer revealed that our approach has excellent power with a low false discovery rate. In nearly all scenarios we tested, JAFFA outperformed other methods for identifying fusions. The only exception was on 50bp paired-end reads, where SOAPfuse had the best performance. When we examined the optimal sequencing read layout and length for fusion detection, we found that JAFFA was the most sensitive on 100bp pair-end reads compared with any other scenario or tool.

The pipeline we have presented is customizable, such that component programs, for example the assembler or aligner, can be easily swapped to current state-of-the-art software. Known fusions that were missed by JAFFA on 50bp reads were lost during the assembly stage. Transcriptome assembly is a relatively recent development and is still maturing. Hence there is potential for JAFFA to produce even better fusion detection sensitivity on short reads in the future.

## Materials and methods

Each stage of the JAFFA pipeline is shown in Figure 1, Additional File 1: Supplementary figure 1, and described below. The pipeline commences by unzipping reads using Trimmomatic [43]. By default, JAFFA does not trim reads but this option is available.

### Preliminary Read Filtering

To aid in computation efficiency, JAFFA begins by filtering out any reads that map to intronic, intergenic or mitochondrial sequence in the genome. This is achieved through a two-step process. Initially all read pairs that map concordantly to the reference transcriptome will be retained. Those that do not map, will move to the second step, where they will be mapped to a version of the human genome, hg19, with exonic sequence masked out. Any read pairs that fail to map concordantly will be retained and merged with those from the initial step. Approximately, 70-95% of reads pass this filter.

### Assemble Reads

Short reads were *de novo* assembled using Velvet version 1.2.10 and Oases version 0.2.08 with k-mer lengths of 19, 23, 27, 31 and 35. We required Oases to output contigs with 100 bases or more. Other settings were default.

### Remove Duplicates

BBMap version 33.41 (http://bbmap.sourceforge.net) was used to remove duplicate reads and convert the fastq reads to fasta format.

### Select reads that do not map to known transcripts

In the case of the Direct mode, reads were mapped as single-end to sequences from GENCODE version 19. We used bowtie2 with the option “-k1--un” for the alignment. For the Hybrid mode, we mapped reads to the GENCODE transcriptome, then took the reads that did not map and attempted to map these to the *de novo* assembled transcriptome. The same bowtie2 settings as above were used.

### Align contigs/reads to known transcripts

We used BLAT [15] to align transcript sequences. When aligning to the transcriptome, we required 98% sequence identity over more than 30 bases, with no intronic gaps, “-minIdentity=98 -minScore=30 -maxIntron=0”. A tile size of 18 was used to improve computational speed, “-tileSize=18”, for the assembly mode, or for reads longer than 100bp, otherwise a tile size of 15 was used to improve sensitivity. These BLAT options are the default in the JAFFA pipeline.

### Select contigs/reads that match multiple genes

We first did a loose selection step to identify which tumour sequences aligned to multiple reference transcripts. The two (or more) reference transcripts were required to be separated by 1kb in the genome by default. Following this we calculated the number of bases that the reference transcripts had in common at the breakpoint If two genes contained the same sequence over a length that was more than the minimum assembly k-mer length (19 bases), a false chimera may be reported. We controlled for this by only selecting fusion candidates with 13 bases or less of sequence in common between the reference genes (Additional File 1: Supplementary figure 2). This step was implemented as an R script

### Counting reads and pairs spanning breakpoints

We counted the number of spanning reads and spanning pairs across the breakpoint. Spanning reads were defined as reads that lay across the breakpoint. Spanning pairs were defined as pairs in which the reads of each pair, lay in their entirety, on opposite sides of the breakpoint. This calculation was performed differently depending on whether the reads were assembled or not. For assembled reads, the reads were mapped back to the candidate *de novo* transcript sequences using bowtie2 with the alignment flags of *“-k1--no-unal--no-mixed--no-discordant”*. Spanning reads were required to have 15 base pairs of flanking sequence either side of the break. For the direct mode, spanning pairs were calculated by mapping reads to the reference transcriptome and searching for discordantly aligned pairs, consistent with the predicted fusion. Each fusion candidate in Direct mode was initially assigned one spanning read (i.e. since the sequence for which the candidate was identified was itself a read). Therefore in this mode, the minimum flanking sequence was 30bp, the minimum to identify a fusion. When multiple reads or contigs predicted the same breakpoint the read support was aggregated.

### Aligning candidate contigs/reads to the genome

We aligned the candidate fusion sequences to the human reference genome (hgl9) using BLAT with default options.

### Check genomic gap, frame and classify candidates

The genomic coordinates of each breakpoint were found and the genomic gap size calculated. In some cases, the gap was very small (less than 10kb) indicating that the candidate was likely to be a false positive, generally due to families of genes with similar sequence or repeated sequence in the genome. These candidates were discarded. Candidates between adjacent genes can also be reported due to run-through transcription or unannotated splicing. We tried to distinguish these scenarios from genuine fusions with small gaps, by looking for evidence of a genomic rearrangement or inversion, based on the direction of the *de novo* transcript with respect to the genome. If no such evidence was found and the gap was less than 200kb the fusion was flagged as a “PotentialRegularTranscript” (not reported by default). Next we determined whether the breakpoints lay on known exon-exon boundaries, as would be expected if the fusion occurred within intronic DNA and the exon structure was preserved. If it did, we checked whether the fusions were in-frame, using the most common frame of the gene’s isoforms. Finally, we grouped candidates that predicted the same genomic breakpoint, aggregated read counts and selected the sequence with the most spanning reads as a representative. For each candidate that was identified by JAFFA we use the spanning reads, spanning pairs, whether the transcriptional breakpoint aligned with exon boundaries and genomic gap to classify then rank the candidates, as described in Results.

### Combine multi-sample results

The pipeline described above was executed in parallel for each sample in a dataset. As a final step, we merged the results from all samples, outputting a table of results and candidate fusion sequences.

### Reference data

The reference transcriptome sequences (GENCODE version 19), exon structure information and human genome version hgl9 were downloaded from UCSC. The reference transcriptomic data is provided with the JAFFA package.

## Dataset Availability

The Edgren and ENCODE dataset can be found on SRA under accession numbers SRP003186 and SRR534293 respectively. The PacBio data can be found at http://blog.pacificbiosciences.com/2013/12/data-release-human-mcf-7-transcriptome.html. The FusionMap simulation is available with the FusionMap software at http://www.arrayserver.com/wiki/index.php?title=FusionMap. The BEERS simulation is available from the authors by request. The gliomas dataset was created from SRP027383. The 13 SRA samples are listed in Additional File 4. Downsampling was non-random, with the first *n* reads being taken from the fastq file using *“head”*. An example is provided in Additional File 5.

## Fusion Finder Comparison

TopHat-Fusion 2.0.13 was run with the parameters similar to those specified on its example website for analyzing the Edgren dataset. Specifically, using the tophat options “--fusion-search --keep-fasta-order --bowtie1 --no-coverage-search --max-intron-length 100000 --fusion-min-dist 100000 --fusion-anchor-length 13 --fusion-ignore-chromosomes chrM,chrUn_gl000220” and the tophatfusion-post options “--num-fusion-reads 1 --num-fusion-pairs 2 --num-fusion-both 5”. For single-end reads we set “--num-fusion-pairs 0”. The insert size (-r) and standard deviation (--mate-std-dev) were modified to match each dataset. For the Edgren dataset we used the values advised on the TopHat-Fusion website. For the FusionMap, BEERS, ENCODE and gliomas datasets we used an insert size of 8, 100, 50 and 0 respectively and a standard deviation of 20, 100, 50 and 100 respectively. When the ENCODE reads were trimmed to 50bp, we increased the insert size to 150. JAFFA 1.04, DeFuse 0.6.2, SOAPfuse 1.26 and FusionCatcher 0.99.3d were all run with default settings. For deFuse, we used the results file that had been thresholded on probability. For all tools, samples within a dataset (e.g. Edgren) were run individually and not pooled. A shell script to reproduce the results from JAFFA is provided as Additional File 5. For the analysis of sensitivity and specificity, we only counted fusion gene pairs with multiple breakpoints once. True positives were identified by their gene name. Any order of gene names was accepted. Different gene aliases were also considered.

## Authors’ contributions

NMD, IJM and AO conceived of ideas in this manuscript. NMD wrote all the code and performed all the analysis. NMD, IJM and AO wrote the manuscript.

## Description of additional files

The following additional data are available with the online version of this paper.

File name: Additional file 1
File format: DOCX
Title of data: Supplementary Figures and Tables
Description of data: Additional file 1 includes eleven supporting figures and two supporting tables. A description of each is given within the file.

File name: Additional file 2
File format: CSV
Title of data: Performance of four transcriptome assemblers on the Edgren dataset
Description of data: A table of which true positive breakpoint sequences were assembled by Trinity, Oases, TransABySS and SOAPdenovo-Trans on the Edgren dataset. Oases assembled the highest number of true positive breakpoints with 31.

File name: Additional file 3
File format: XLSX
Title of data: Fusion genes in the BT-474, SK-BR-3, KPL-4 and MCF-7 cell lines Description of data: A list of the true positive fusion genes used in the validation of JAFFA on the Edgren and ENCODE dataset, along with a list of the probable true positives, and the fusion calls from JAFFA.

File name: Additional file 4
File format: XLSX
Title of data: Fusion genes in the glioma dataset.
Description of data: A list of the true positive fusion genes, probable true positives and results from JAFFA for the gliomas dataset.

File name: Additional file 5
File format: sh
Title of data: JAFFA commands
Description of data: This script provides commands to reproduce the results from JAFFA shown in the manuscript.

## Acknowledgements

We would like to acknowledge Simon Sadedin for his help with Bpipe, Aliaksei Holik and Alex Gout for their feedback as pilot users of the software, Katrina Bell and Maria Doyle for their feedback on this manscript, and Lorenza Mittempergher, Chong Sun and Astrid Bosma for suggestions on the output of JAFFA. This work was made possible through Victorian State Government Operational Infrastructure Support and Australian Government NHMRC IRIISS. This work was supported by the National Health and Medical Research Council, Australia (Career Development Fellowship 1051481 to AO, Project grant 1051402 to AO, Program Grant 1016647 to IJM, and Fellowship 575581 to IJM).

